# Assessment of CAR-T mediated and targeted cytotoxicity in 3D microfluidic TBNC co-culture models for combination therapy

**DOI:** 10.1101/2021.09.14.458168

**Authors:** K. Paterson, S. Paterson, T. Mulholland, S. B. Coffelt, M. Zagnoni

## Abstract

Chimeric antigen receptor (CAR)-T cell therapy is efficacious against many hematological malignancies; however, their therapeutic application to treat solid tumors presents further challenges. A better understanding of how the solid tumor microenvironment (TME) impacts CAR-T anti-tumor effects would enable the selection of effective combination therapies to decipher the optimal course of treatment for patients and to better engineer CAR-Ts. Classical 2D *in vitro* models do not provide sufficient recapitulation of the native human TME, and *in vivo* models, such as patient-derived xenografts, are costly, complex and labor intensive. Here, we present a novel 3D, miniaturized assay for the evaluation of EGFR-targeted CAR-T cell cytotoxicity and specificity on tumor-stroma triple-negative breast cancer models in microfluidic devices. CAR-T cells were shown to home towards EGFR-expressing cancer cells to elicit a cytotoxic effect, whilst leaving low EGFR-expressing fibroblasts viable, an effect which was enhanced through combination anti-PD-L1 therapy and carboplatin chemotherapy. Hence, we propose this proof-of-concept immunoassay as a future preclinical screening tool for the development of novel immunotherapeutics and for use in personalized medicine.

## Introduction

Epithelial malignancies account for up to 90% of all cancers and exhibit an array of evasive mechanisms to escape immune surveillance.^1, 2^ Triple-negative breast cancer (TNBC) makes up to 20% of breast cancer cases, is highly aggressive and lacks successful therapeutic options.^3, 4^ TNBC is characterized by an absence of estrogen receptor (ER), progesterone receptor (PR) and human epidermal growth factor receptor 2 (HER2).^4^ The epidermal growth factor receptor (EGFR) is expressed in the majority of cancer cells and also in many types of TNBC and is a promising target for the development of novel immunotherapies.^3, 5^ Programmed cell death protein 1 (PD-1) and programmed cell death ligand 1 (PD-L1) are responsible for the inhibition of immune responses and modulation of T cell activity.^6^ The PD-1/PD-L1 pathway plays an important role in tumor escape of immune surveillance and can lessen the effectiveness of anti-cancer therapies.^6^ Combination anti-PD-L1 and chemotherapy treatment for TNBC is more efficacious than individual monotherapies in terms of progression-free and overall survival.^7^ Carboplatin chemotherapy is commonly used in the treatment of TNBC^8^, with several trials underway to investigate various combination regimes including carboplatin and PD-1/PD-L1 inhibitors.^9, 10^ Both animal and ex vivo studies have shown that combination treatment caused a greater reduction in tumor volume and extended survival in comparison to the individual monotherapies.^7, 8^

Chimeric antigen receptor (CAR)-T cell immunotherapies are remarkably successful in the treatment of hematological malignancies and focus has now shifted to harnessing this technology towards tackling solid tumors.^1^ As examples, the most targeted antigens in CAR-T clinical trials in patients with epithelial malignancies have been EGFR, HER2 and natural killer group 2D (NKG2D)-ligands.^11^ However, there are several factors impairing widespread application of CAR-T therapy. CAR-T cell production is associated with high manufacturing costs due to the autologous acquisition process from patients.^12^ Off-target toxicity can trigger serious or even life-threatening therapy response.^1, 13 14^ Specific to solid tumors, CAR-T cell infiltration is hindered by the immunosuppressive tumor microenvironment (TME). The solid tumor milieu comprises a variety of cell types releasing an assortment of cytokines, chemokines, growth factors and immune checkpoint molecules that aid tumor growth and lessen the effectiveness of CAR-T therapy.^15^ As an example, cancer associated fibroblasts (CAFs) are known to inhibit T cell access to tumor cells.^1, 16, 17^ To enhance CAR-T tumor infiltration, chemotherapy or radiotherapy can be used.^1^ Several trials have investigated combining chemotherapy and immune checkpoint blockade to create a more hospitable immune microenvironment for CAR-T cells.^9, 18, 19^ Due to the variety of mechanisms implicated in immunosuppression, there are many combination treatments and CAR-T designs that could be trialled.^1, 15^ Therefore, predictive preclinical models that better mimic the challenges associated with tackling solid tumors are needed for screening such variations. As a solution, miniaturized 3D assays could provide a cost-effective approach for studying CAR-T efficacy and off-target quantification by performing advanced mechanistic studies.

*In vitro* CAR-T assays are typically conducted in 2D, but most recently these have increasingly been developed in 3D with low-adhesion well plates or hanging drop techniques, ^3 5 20 21 22 23 24 25^ using a wide range of readouts, such as kinetic, cytokine release, viability and activation state analysis. However, immunoassays that incorporate tumor spheroid and stromal co-cultures are not widely used.

Miniaturized immunoassays have been developed using microfluidic technology,^26 27 28 29 30 31 32^ but their application is still limited in relation to CAR-T studies.^33^ As examples, Ando *et al*. established a microfluidic assay to study the effect of hypoxic condition to CAR-T cell behaviour.^34^ Pavesi *et al*. studied T cell efficacy in an inflammatory and hypoxic microenvironment.^35^ Notably, 2D assays showed significantly greater killing by T cells in comparison to 3D microfluidic studies, emphasizing the importance of 3D models during *in vitro* modelling. With respect to off-chip CAR-T screening approaches, microfluidic technology can be effective in increasing the complexity of the assays (such as for combination therapy studies) whist miniaturizing the assessment of CAR-T therapeutic strategies, thus decreasing assay costs and time to results.

In this paper, we have developed a novel microfluidic method to assess CAR-T cell-mediated cytotoxicity and off-target identification on multiple TNBC cancer-stroma co-culture spheroids. This work is the first example considering how combination treatments, consisting of carboplatin chemotherapy, anti-PD-L1 therapy and CAR-T therapy, influence CAR-T killing efficiency in 3D. EGFR-targeted CAR-T cells were selected as the EGF is a receptor over-expressed in many solid tumor types.^36^ 3D co-cultures of cancer cell with high EGFR expression levels and normal healthy fibroblast with low EGRF expression levels were used to assess CAR-T killing specificity. 3D co-cultures of cancer cells with CAFs (largely neglected in *in vitro* CAR-T models^37^) were also studied, as many solid tumors can be composed of this cell type. CAFs are an important feature of the TME and are implicated in the outcomes of many therapies.^17, 33, 38^ These models were then used to investigate the CAR-T mediated cytotoxicity in combination studies that mimic clinical TNBC regiments.^19, 39^ Image analysis provided quantification of cell-mediated cytotoxicity in relation to therapy-induced cell expression levels and effector-target ratio. Results from microfluidic experiments showed how CAR-T killing and targeting of cancer cells was enhanced in combination studies with respect to controls.

This proof-of-concept work offers evidence of how the microfluidic platform and protocols can provide powerful and miniaturized *in vitro* assays to assess CAR-T cell therapies in preclinical settings. The methods can also be adapted for a wide range of adoptive cell therapy studies.

## Experimental section

### Cell monolayer culture

All cells were incubated at 37 °C and 5% CO2 in a humidified incubator. MDA-MB-468 were kindly provided by Prof E. Piletska (Department of Chemistry, University of Leicester). The human mammary CAFs, hTERT immortalized and GFP positive, were generated by Kojima et al.^40^ and were kindly provided by Prof Sara Zanivan (CRUK Beatson Institute, Glasgow). MDA-MB-468 and CAF were both cultured in DMEM (ThermoFisher) supplemented with 1% penicillin/streptomycin (10000 units/ mL, ThermoFisher) and 10% heat-inactivated FBS (ThermoFisher). Normal human lung fibroblasts (NHLF) were purchased from Lonza and cultured in ready to use PrimaPure™ Fibroblast Growth Medium (AMS Biotechnology (Europe) Limited). EGFR-TM28-GITR-CD3z CAR-T cells and CAR-T cell media containing FBS were both obtained from AMS Biotechnology (Europe) Limited. Upon thawing, CAR-T cells were centrifuged and their supernatant discarded. CAR-T cells were then resuspended in pre-warmed media and incubated overnight prior to use the following day.

### 3D cell culture in microfluidic devices

Cells were cultured in *OC^3D^ Single* microfluidic devices (ScreenIn3D Ltd, UK), consisting of an array of 24 independent culture chambers (Figure 1A). Each chamber hosts 25 co-culture spheroids within ultralow-adhesion microwells (250 × 250 × 200 μm^3^), which are fluidically addressable by a microfluidic channel connected by two open wells. Devices were fabricated as previously described.^41^ Devices were washed using phosphate-buffered saline (PBS) and stored at 37 °C and 5% CO_2_ in a humidified incubator prior to cell seeding. Cells were seeded into devices (3-7μL at a concentration of 2-7.5× 10^6^ cells/mL) to form spheroids and medium replenished every 24 to 48 hours. MDA-MB-468 and stromal cells, CAF or NHLF, were seeded into devices both as 3D mono and co-cultures.

**Figure 1.**
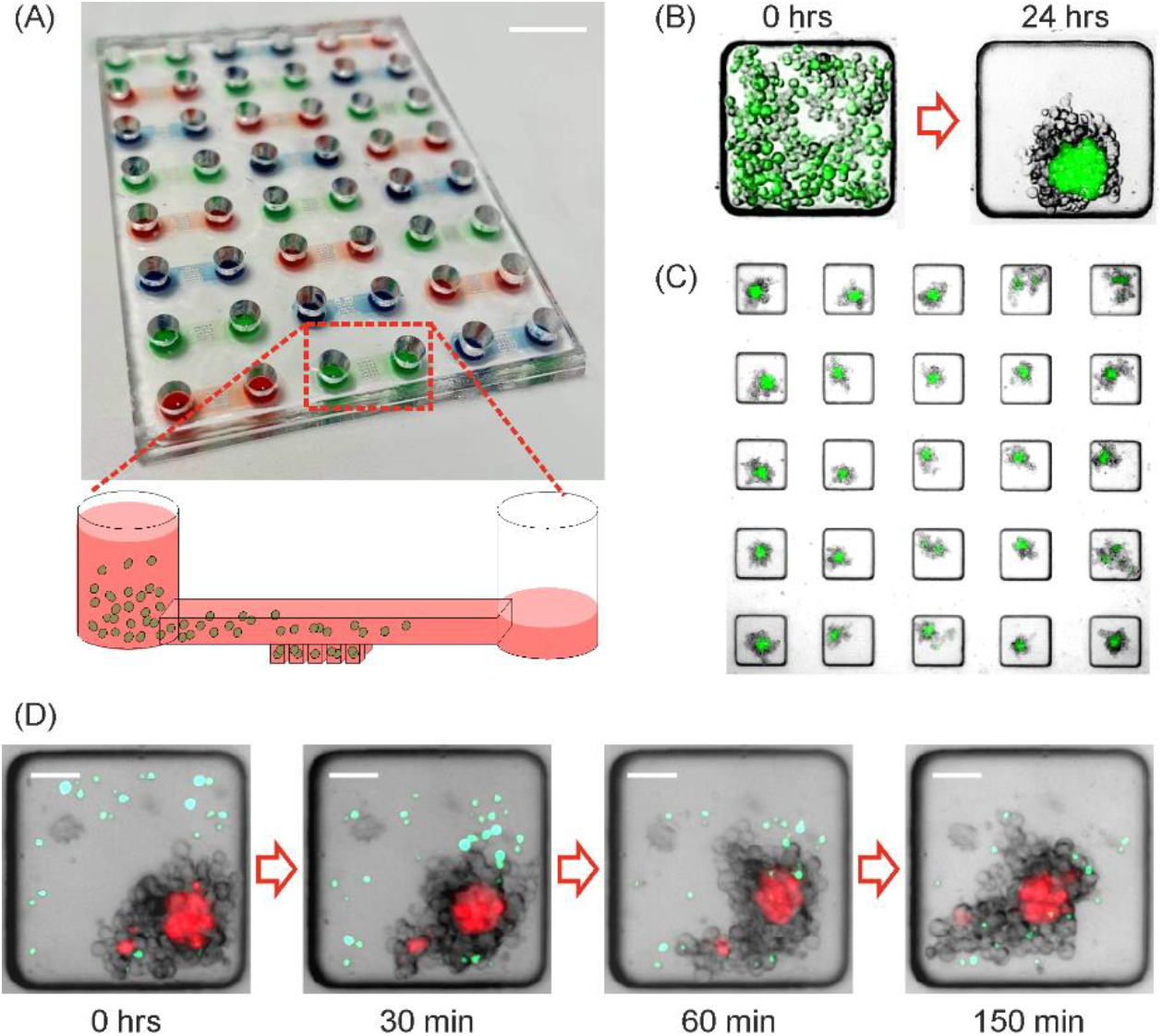
Microfluidic tumor-stromal 3D co-culture for CAR-T studies. (A) Image of the OC^3D^ Single microfluidic device used (ScreenIn3D Ltd, UK) with schematic showing principle of cell seeding. A single-cell suspension flows through a microchannel and cells sediment into microwell traps below the microchannel level. Scalebar = 10 mm. (B) Brightfield and fluorescence images showing spheroid formation within 24h following seeding of MDA-MB-468 (unlabeled) and CAF (green). (C) Brightfield and fluorescence image of an array of microwell traps, allowing the culture of 25 spheroids. (D) Brightfield and fluorescence images from a time-lapse experiment showing CAR-T cells (green) homing and interacting with MDA-MB-468 (unlabeled) and NHLF (red) co-culture spheroids. CAR-T cells were injected immediately prior to beginning time-lapse recording. Scalebar = 50 μm.

### CAR-T and Combination therapy assay

Carboplatin stock solution was diluted in cell culture media to the desired concentration for experiments (12.5-200μM) and added for 24h to devices once cells had aggregated to form spheroids on day 1. On day 2 of culture, anti-PD-L1 antibodies (329701, Biolegend) were injected into devices after a 1:100 dilution in culture media. On day 3 of culture, all media was removed from devices and CAR-T cells injected (3μl at a concentration of 2.5-10×10^6^/mL) and incubated for up to 72h with the spheroids. Prior to injection, cells were fluorescently labelled using dyes that were freshly prepared from CellTrace™ Far Red, Bue or CSFE Proliferation Kits (Thermofisher), depending on the experimental set-up. 20uL of DMSO (Sigma) was added to a CellTrace stock vial. Cells in suspension were centrifuged and resuspended in pre-warmed PBS buffer (PBS/2%FBS) to a maximum concentration of 1×10^6^ cells/mL. 1μL of CellTrace dye per mL of cell suspension was then added and cells incubated for 20 minutes at 37°C. After the incubation period, ice-cold quenching solution at 5 times the volume of the cell-CellTrace staining solution was added. Cells were then centrifuged and the supernatant removed. Labelled cells were resuspended in the desired volume for use in experiments. Anti-EGFR antibody (ab231, abcam) was used at a 1:200 dilution in cell culture media for EGFR blocking experiments. Anti-EGFR was added to devices on day 1 of culture for 24h before removal of all media from devices and addition of CAR-T cells for 72h. Every experiment was performed in triplicate, with at least 50 spheroids analyzed per condition tested.

### Cell viability

To determine spheroid viability, 5mg/mL fluorescein diacetate (FDA, Sigma) and 2mg/mL propidium iodide (PI, Sigma) stock solutions were freshly prepared in acetone and PBS respectively. FDA and PI were diluted in media to final concentrations of 8μg/mL and 20μg/mL and added to devices. Cells were incubated in the staining solution for 30 minutes at 37°C. PBS was subsequently used to wash out excess staining solution and was refreshed prior to imaging. After staining experiments were terminated.

### Immunofluorescence

For the quantification of EGFR and PD-L1 expression, solutions of PBSB, PBS containing 0.1% BSA (Sigma), and PBSBDT blocking solution, PBS containing 0.5% Triton-X (Fisher), 1% DMSO, 1% BSA and 1% FBS were freshly prepared on the day prior to fixing. All medium was removed from devices before washing with PBSB and incubation on ice for 30 minutes. PBSB was removed and 4% paraformaldehyde (PFA) added to devices for another 30 minutes.

PFA was removed and devices washed again with PBSB prior to incubation with PBSBDT for 1 hour. The blocking solution was removed and primary antibodies for either EGFR, recombinant anti-EGFR antibody (ab32198, abcam), or PD-L1, purified anti-human CD274 (B7-H1, PD-L1) antibody (329701, Biolegend), were added at a 1:100 dilution for 24-48h and stored at 4°C. Devices were washed with PBSB and incubated at room temperature for 10 minutes before the addition of secondary antibodies in a 1:200 ratio. Goat anti-Mouse Alexa Fluor 633 Secondary Antibody (Thermofisher) was added for PD-L1 quantification and Goat anti-Rabbit Alexa Fluor 633 Secondary Antibody (Thermofisher) was added for EGFR quantification. Secondary antibodies were incubated in devices for 2 hours at room temperature before washing steps and imaging.

### Statistical analysis

Statistical analysis was carried out with GraphPad Prism 8.1.2 (GraphPad Software, Inc., San Diego, CA). All data is presented as mean ± standard deviation, unless otherwise stated, using bar graphs or scatter plots. T-tests were used for the comparison of two variables. For comparison of multiple variables, two-way analysis of variance (ANOVA) with Bonferroni multiple comparison tests were conducted. Differences between groups were considered to be significant at a P value of <0.05. For data analysis, spheroids were selected only if presenting on day 2 of culture a radius of 60 ± 15 μm for cancer cell monocultures, a radius of 25 ± 10 μm for fibroblast monocultures and a radius of 40 ± 10 μm for cancer cell and fibroblast co-cultures. At least 50 spheroids were considered for statistical analysis per condition tested.

### Microscopy setup and image analysis

Cultures were imaged via bright-field and epifluorescence microscopy using an inverted microscope (Axio Observer Z1, Zeiss) connected to an Orca Flash 4.0 camera (Hamamatsu). Images were collected daily and before and after drug treatment and CAR-T injection. For real-time imaging experiments, devices were kept in a temperature and humidity controlled stage top incubator (OKOLAB, Italy) for automated imaging overnight at 37°C at 94% relative humidity with a gas flow rate of 0.1 l/min.

Image analysis was performed using ZEN Blue and Fiji to measure spheroid area and to count CAR-T cells. Co-localization of cancer cell with PI signal was quantified by extracting individual TIFF images from each wavelength channels using Fiji and normalizing them to the same threshold range. Subsequently, TIFF images were converted to an 8-bit binary image for area and cell number quantification. The area of CAF (obtained from GFP channel) and area of CAR-T cells (obtained from CellTrace Far Red channel) were subtracted from the PI signal area, enabling estimation of co-localised area of PI signal with unlabelled cancer cells, plotted as a percentage of total PI area.

## Results

To establish a co-culture system for EGFR-specific CAR-T-mediated cytotoxicity, we selected MDA-MB-468 breast cancer cells for their reportedly high expression of EGFR.^42–44^ We also chose two stromal cell types with low level expression of the EGFR receptor: immortalized cancer-associated fibroblasts (CAF) from a breast tumor and normal human lung fibroblasts (NHLF). All three cell types were tested for EGFR expression. Results were consistent with previous reports, showing that MDA-MB-468 expression of EGFR was approximately 4 times greater than in CAF and 5 times greater than in NHLF (Figure S1 in SI), providing positive and negative targets of EGFR-specific CAR-T cells.

### Assessment of EGFR Specificity of CAR-T in 2D

Tests were performed initially in 2D to assess the cytotoxic activity of CAR-T cells towards the EGFR target. For this, monocultures of MDA-MDB-468 were established in 96-well plates (Figure S1 in SI). Assays investigating the effect of varying effector to target (E:T) ratio, testing 1:2, 1:1 and 5:1 E:T ratios, showed that the number of live MDA-MB-468 cells decreased with increasing E:T ratio after 72h incubation with CAR-T cells. The number of CAR-T cells remained constant throughput the duration of the assay. Even at an unfavorable 1:2 E:T ratio, CAR-T elicited a significant cytotoxicity effect on cancer cells.

### 3D co-culture models

Therefore, considering these results (Figure S1C in SI) and aiming to validate cost-effective 3D assays that minimize usage of CAR-T material, microfluidic 3D experiments were designed to obtain a 1:10 E:T to visualize CAR-T interaction with the cancer spheroid model (Figure 1, quantification of seeded CAR-T cells in microwells confirmed this condition, not shown). The high specificity of these CAR-T cells meant that E:T ratios lower than typical values for *in vitro* assays could be used to produce statistically significant cytotoxic results. Throughout the development of this assay, protocols were refined to use as few as 7,500 T cells/injection in each culture chamber, achieving a mostly uniform distribution of CAR-Ts in the 25 microwells (approx. 20 CAR-Ts per 200 cancer cells). This outcome is evidence of the miniaturization capabilities of the platform and its potential benefit for future assays with patient biopsy tissue.

Having validated high EGFR expression in breast cancer cells and low EGFR expression in fibroblasts, MDA-MB-468 and the two fibroblasts were seeded at a 1:1 ratio in *OC^3D^ Single* microfluidic devices, hosting 24 array chambers (Figure 1A). Due to the ultra-low adhesion conditions, cells aggregated into a spheroid within 24 hours (Figure 1B). Each microfluidic chamber enable the simultaneous formation and culture of 25 spheroids per condition tested, with radius ranging 45-71μm for cancer cell monoculture spheroids, 17-28μm for fibroblast monoculture spheroids and 32-50μm for cancer cell and fibroblast co-culture spheroids (Figure 1C). After spheroid formation, CAR-T cells were injected for co-culture with the spheroids. CAR-T cells migrated towards the spheroids immediately after injection and, within 3 hours, each CAR-T cell could be seen interacting with the spheroid mass (Figure 1D) (MovieS1 in SI).

### Assessment of CAR-T targeting and cytotoxicity

3D cultures were set up in microfluidic devices to test both the CAR-T targeting specificity and their mediated cytotoxicity on MDA-MB-468 and fibroblast spheroids, both alone and as co-cultures (Figure 2).

**Figure 2.**
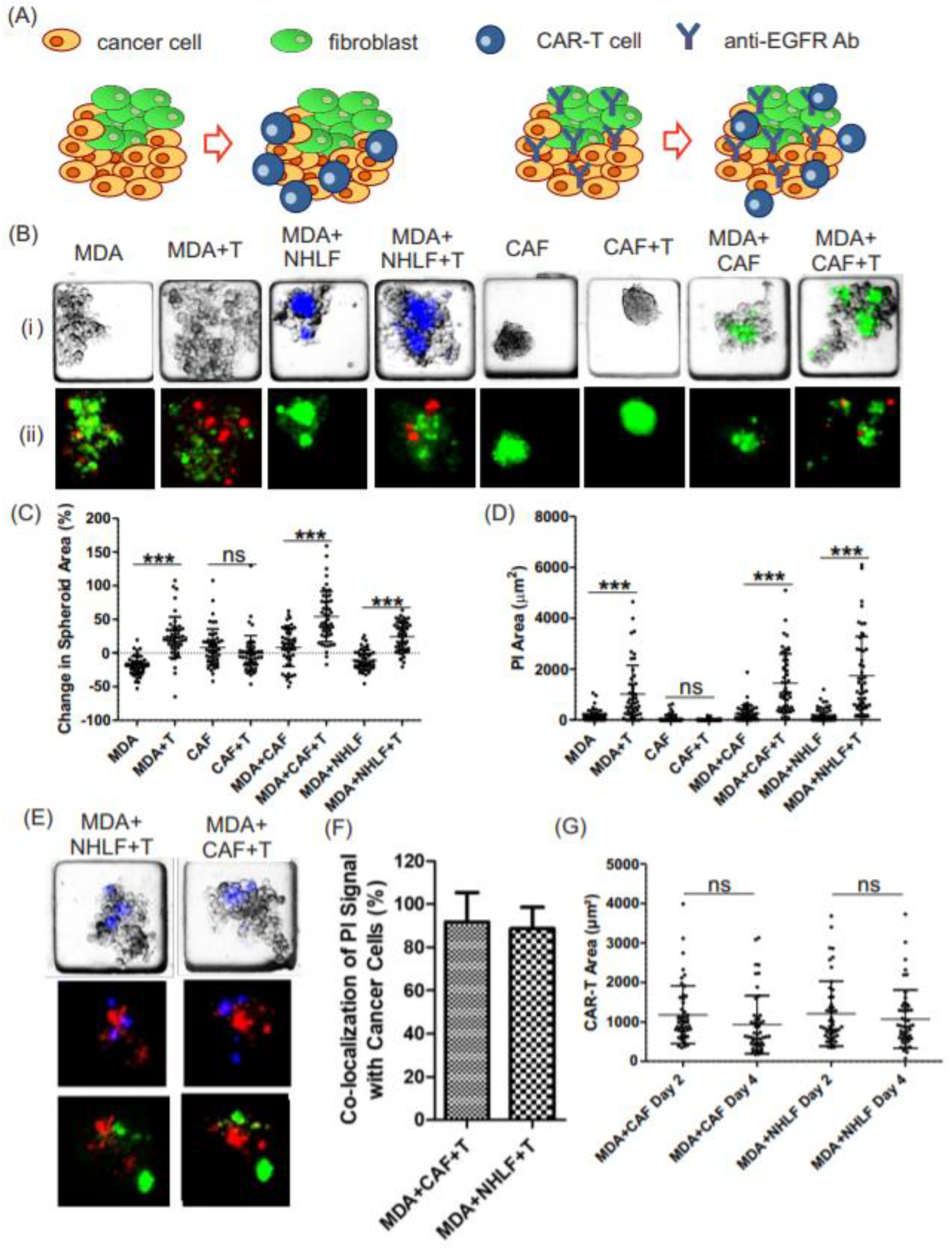
CAR-T targeted specificity and cytotoxicity in mono- and co-cultures. (A) Schematic of experimental set-up showing cancer cell (yellow) and fibroblast (green) co-culture spheroids treated with CAR-T cells (blue) and anti-EGFR antibodies. (B) Representative images of spheroids from MDA-MB-468 and CAF monocultures and MDA-MB-468 spheroids in co-culture with CAF (green) and NHLF (blue) in different culture conditions after 72h CAR-T incubation. (i) Brightfield and fluorescence images. (ii) Fluorescence images obtained after staining with FDA (green) and PI (red). For CAF mono- and co-cultures only PI signal is shown (red). (C) Plot of percentage change in spheroid area from day 2 to day 5 of culture (n=50). (D) Plot of PI area signal (n=50). (E) Representative brightfield and fluorescence images of MDA-MB-468 (unlabeled) in co-culture with NHLF/CAF (green) after 72h of CAR-T (blue) incubation and stained with PI (red). (F) Plot of percentage of PI signal co-localized only with unlabeled cancer cells (n=50). (G) Plot of area representative of CAR-T cell coverage in microfluidic devices over time, n=50. MDA=MDA-MB-468, T= CAR-T cells.

Spheroids aggregated for 24h prior to CAR-T cell injection, obtaining a 1:10 CAR-T to cancer cell ratio. CAR-T cells were incubated for 72 hours prior to imaging and viability staining. Two outcomes resulted from this assay: first, CAR-T induced disaggregation of MDA-MB-468 spheroids over the first few hours of incubation, irrespectively of whether monoculture and co-culture spheroids were used. No significant change to the area of CAF monoculture spheroids incubated with CAR-T cells was observed (Figure 2C). Lastly, an elevated marker of cell death (intensity from PI signal) was obtained for MDA-MB-468 spheroid monocultures and MDA-MB-468-CAF and MDA-MB-468-NHLF spheroid co-cultures with respect to controls after 72 hours CAR-T incubation (Figure 2D). No significant change in cell death was detected for CAF monoculture spheroids after 72h CAR-T treatment. This data suggests that CAR-T cells predominantly targeted EGFR high-expressing cells. To further confirm this finding, 2D image analysis was performed to quantify PI signal co-localization with cancer cells, fibroblasts or CAR-T cells (Figure 2E). For NHLF and CAF co-cultures, the mean and standard error of mean values for the percentage of PI signal co-localized to cancer cells was quantified as 88.6 ± 1.4% and 91.7 ± 1.9%, respectively (Figure 2F). Considering the lack of cell death obtained from fibroblast spheroid mono-cultures (both CAF and NHLF, Figure 2D) and that we could perform only 2D image analysis of a 3D cellular structure (thus potentially associating PI signal overlap with stained fibroblasts even if at different height levels in the spheroid), the results are indicative of high level, on-target specificity of the CAR-T cells. It should be noted that during the live/dead staining process, some dead cells could be easily washed out from the microwell traps, presumably due to the high damage produced by the CAR-T cells. Similarly to the results from 2D assays, no change to CAR-T numbers was recorded in device cultures (Figure 2G).

In order to determine whether these cytotoxic effects were produced only by CAR-T cells binding to EGFR, an anti-EGFR antibody was incubated with MDA-MB-468 and CAF spheroid monocultures for 24 hours prior to CAR-T cell injection (Figure S2) and washed out. Cultures were subsequently incubated for 72 hours with CAR-T cells. The EGFR antibodies did not cause any significant change in spheroid area or viability when administered alone with respect to controls (Figure S2C-D in SI). However, when present with spheroid co-cultures and CAR-T cells, they prevented both MDA-MD-468 spheroid disaggregation and a significant increase in cell death (measured by PI signal) (Figure S2C-D in SI). These results confirmed CAR-T cell-mediated cytotoxicity and target specificity through EGFR recognition and that this effect could be neutralized with administration of an anti-EGFR antibody.

### Combination therapy

Having characterized the effects and the efficacy of CAR-T administration alone, further studies were performed to assess the outcomes when CAR-T was combined with other anticancer therapies (Figure 3). Based on clinical TNBC therapy regiments, carboplatin and anti-PD-L1 antibody treatments were selected for use in conjunction with CAR-T therapy. For this, after spheroid formation, cells were exposed to carboplatin for 24h, after which all media was removed and anti-PD-L1 antibodies were incubated with the spheroids into devices for another 24h period. After this, all medium was removed and CAR-T cells added to the devices for 72h.

**Figure 3.**
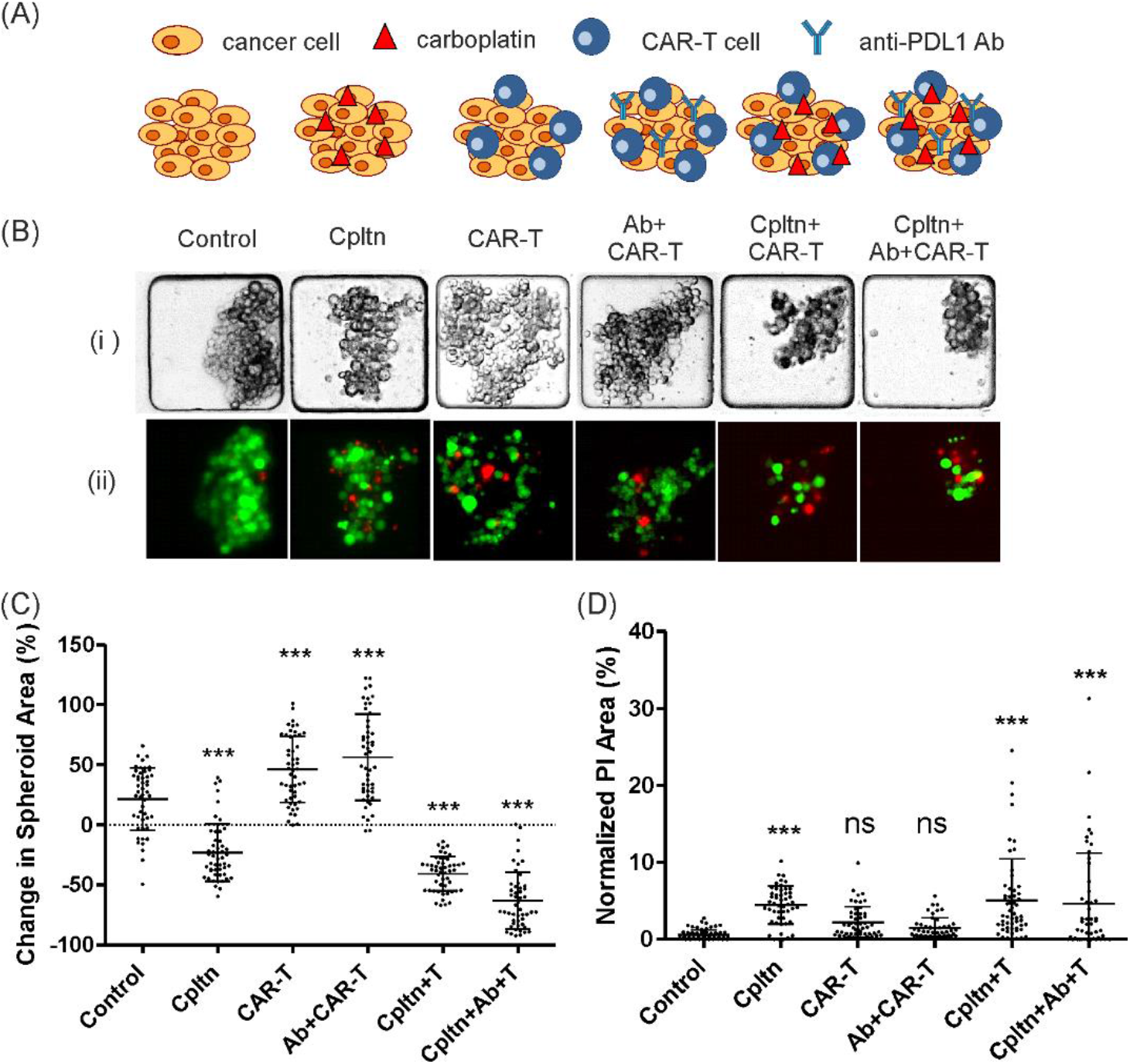
Analysis of the effects produced by combination therapies in MDA-MB-468 spheroid monocultures. (A) Schematic of the different combination therapy modalities used. (B) Representative (i) brightfield and (ii) fluorescence images of MDA-MB-468 monoculture spheroids on day 6, after viability staining with FDA (green) and PI (red). (C) Plot of the percentage change in spheroid area, measured from brightfield images, from day 1 to day 6 (n=50). (D) Plot of the percentage of brightfield area co-localized with PI signal (n=50). Cpltn = carboplatin chemotherapy, Ab = Anti-PD-L1 antibodies, T= CAR-T cells.

Prior to this, MDA-MB-468 spheroid monoculture’s sensitivity to carboplatin was tested to identify the EC50 and then chose the optimum chemotherapeutic dose that would produce a mild chemotoxic effect, maintaining acceptable spheroid viability and aggregation in order to assess any synergistic effects produced by CAR-T cells in all possible conditions. Concentrations ranging from 12.5 to 200μM were tested against MDA-MB-468 spheroids, incubating the drug for 24 hr and terminating the assay on day 6 of culture, mirroring the endpoint used for combination therapy studies. Results showed an EC50 of 28μM (Figure S3B in SI). Brightfield and live spheroid area were found to decrease with increasing carboplatin concentration, whilst PI area and spheroid disaggregation increased with greater doses (Figure S3C-E). Accurate area values were not attainable for spheroids exposed to higher drug concentrations due to extreme disaggregation. The concentration of carboplatin used for combination treatments was selected as 12.5μM, which produced a non-negligible but mild cytotoxic effect (Figure 3).

Results from combination assays on MDA-MB-468 spheroids showed a statistically significant reduction in spheroid areas when chemotherapy was administered alone and in combination with anti-PD-L1 and CAR-T cells (the latter at a greater extent) (Figure 3C). The greatest reduction in cancer cell spheroid area was found to occur when all therapies were combined. A significant increase in the percentage dead area of the spheroid was only recorded for conditions that included carboplatin, given alone or in combinations (Figure 3D). However, as previously noted, many dead cells were washed out from the microwell traps after staining. In the absence of chemotherapy treatment, no statistically significant changes could be detected between CAR-T treatment alone or CAR-T and anti-PD-L1 therapy, suggesting a minor effect of the checkpoint inhibitor in these experiments. Following these studies, the same therapy conditions were also tested against CAF and MDA-MB-468 spheroid co-cultures. (Figure 4).

**Figure 4.**
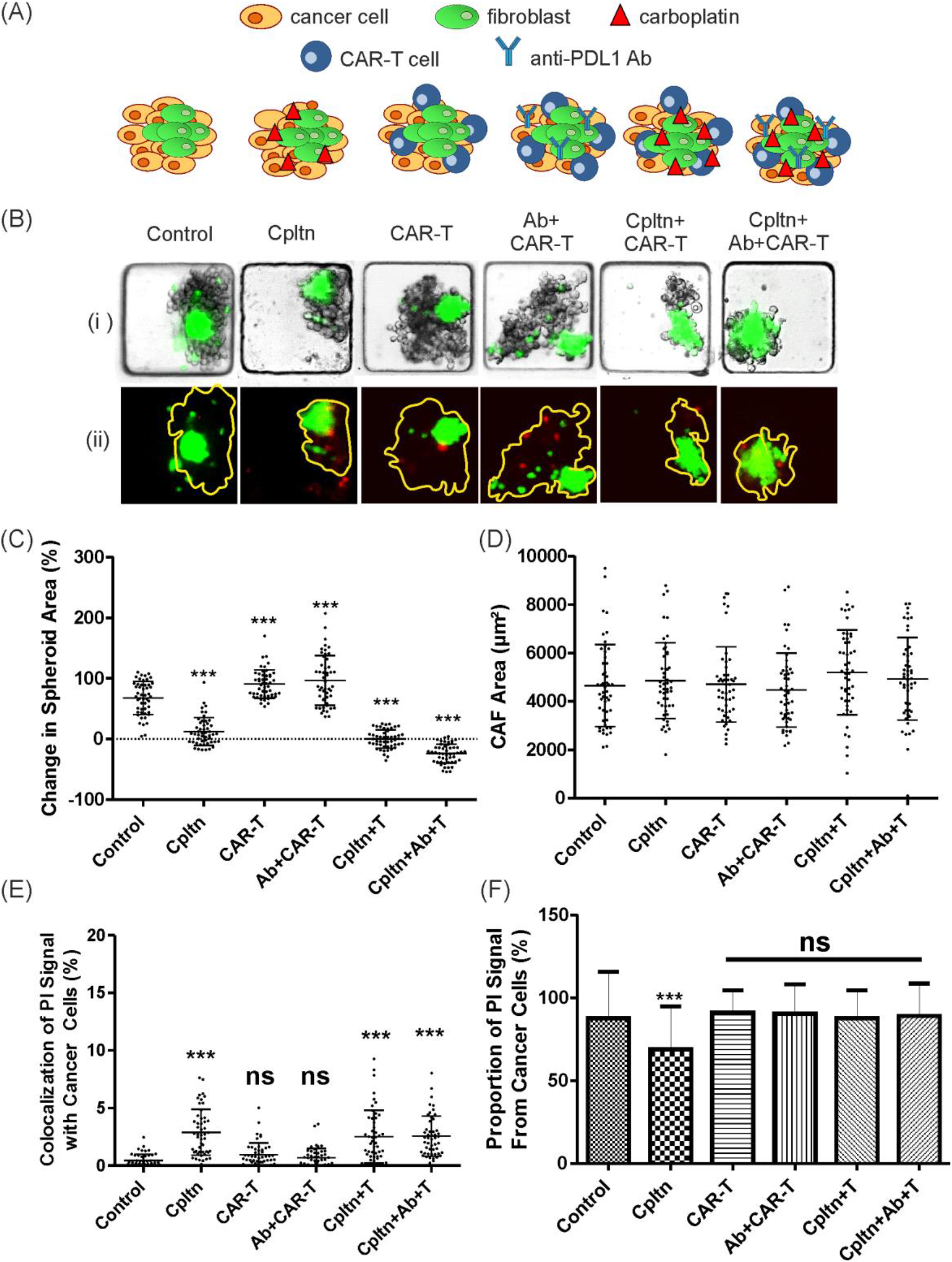
Analysis of the effects produced by combination therapies in MDA-MDB-468 and CAF spheroid co-cultures. (A) Schematic of the different combination therapy modalities used. (B) Representative brightfield and fluorescence images (i) of MDA-MB-468 and CAF (green) co-culture spheroids on day 6, and (ii) after viability staining with PI (red). Yellow outline represents spheroid area. (C) Plot of the percentage change in spheroid area from day 1 to 6, measured from brightfield images (n=50). (D) Plot of CAF spheroid area on day6 of culture extracted from co-culture experiments (n=50). (E) Plot of the percentage of spheroid area co-localized with PI signal (n=50). (F) Plot of percentage of PI signal co-localized only with unlabeled cancer cells (n=30). Cpltn = carboplatin chemotherapy, Ab = Anti-PD-L1 antibodies, T= CAR-T cells.

Consistent with the results obtained from MDA-MDB-468 monoculture conditions, the spheroid area was significantly reduced when chemotherapy was administered alone and in combinations with CAR-T and anti-PD-L1 treatment and CAR-T (Figure 4C). Once again, the largest reduction in spheroid mass was recorded where chemotherapy, anti-PD-L1 and CAR-T were combined. CAFs, expressing low levels of EGFR, were resistant to all combination therapy tested (Figure 4D), showing no statistically significant differences in spheroid shape or size when in co-culture. A statistically significant increase in dead cancer cells was recorded for chemotherapy alone and in combination with CAR-T and anti-PD-L1 treatments (Figure 4E). The combination of anti-PD-L1 treatment with CAR-Ts did not show an increase in cancer cell death, with the chemotherapy treatment being the main discriminant for enhancing the cytotoxic effects of CAR-T cells. Also for these combination studies, the majority of the PI signal (on average 89.63%) originated from cancer cells in comparison control cultures (87.73%) and carboplatin monotherapy (68.94%), demonstrating the on-target specificity of the CAR-T under investigation (Figure 4F).

Some chemotherapies have been reported in the literature to increase tumor cell expression of EGFR^45–47^ and PD-L1^48–50^ for several cancer types. Thus, the levels of EGFR and PD-L1 expression were assessed in the microfluidic for MDA-MB-468 monoculture spheroids after exposure to carboplatin in order to better interpret the results from the 3D assays (Figure 5). Spheroids were treated with carboplatin for 24h after formation and subsequently fixed and stained for PD-L1 and EGFR to quantify expression levels caused by chemotherapy.

**Figure 5.**
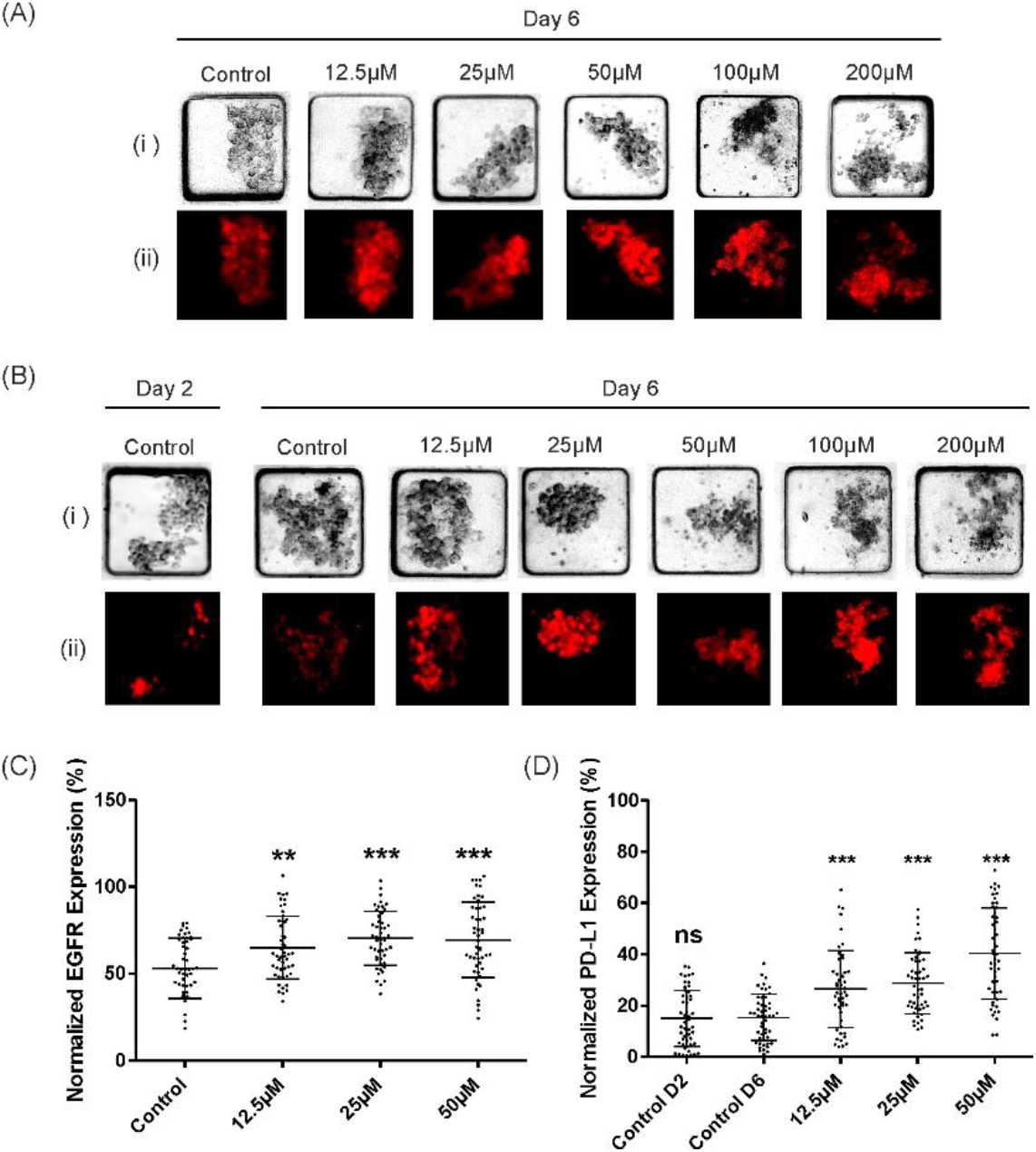
Effect of carboplatin treatment on PD-L1 and EGFR expression levels in MDA-MB-468 spheroids. (A) Representative (i) brightfield and (ii) fluorescent images of MDA-MDB-468 spheroid monocultures stained with anti-EGFR antibody (red) following carboplatin incubation. (B) Representative (i) brightfield and (ii) fluorescent images of MDA-MDB-468 monocultures stained with anti-PD-L1 antibody (red) following carboplatin incubation. (C) Plot of EGFR expression normalized to spheroid area (n=50). (D) Plot of PD-L1 expression normalized to spheroid area on day 6 of culture (n=50). MDA = MDA-MB-468, D2 = Day 2, D6 = Day 6.

Chemotherapy exposure was found to trigger an increase in the expression of EGFR (Figure 5A-B) as well as PD-L1 (Figure 5C-D), for all carboplatin concentration values tested with respect to controls. This response to chemotherapy explained the enhanced CAR-T cell-mediated cytotoxicity observed in the combination therapy conditions, as increased level of EGFR increased the likelihood of CAR-T cell binding. Even if PD-L1 expression levels were increased, this had a secondary (almost negligible) effect on CAR-T efficacy with respect to overexpression of the target. Overall, these results show how a combination of these agents was able to elicit a more powerful anti-cancer response in comparison to monotherapies.

## Discussion

This proof-of-concept work provides an example as to how miniaturization through lab-on-chip technologies can be utilized to augment output data when using limited resources. Due to the high selective toxicity of the CAR-T cells and miniaturized platform used in this work, the E:T ratio could be reduced to far lower than is typical for conventional in vitro studies whilst still eliciting significant cytotoxic effects on the tumor mass. Continuous refinement of cell seeding protocols during this work has resulted in the ability to perform over a hundred 3D CAR-T assays using just 1 million cells. The system can be also adapted to depict a variety of TME conditions in 3D, favoring the development of mechanistic studies for preclinical *in vitro* assessment of CAR-T safety, efficacy and off-target effects prior to *in vivo* studies. Traditional models in 2D can enable CAR-T cell interaction with cancer cells through gravitational forces. In this model, CAR-T cells are able to move freely within microwells, allowing their homing abilities to be more faithfully recapitulated and observed. The microfluidic platform offers the potential to expand these studies to other adoptive cell therapy strategies for solid tumors, providing a flexible platform for precision medicine using patient derived tissue.^41^ Further insights into immune cell function and cytotoxicity could be provided through alternative readouts, such as measuring cytokines or chemokines present in the device culture media. The platform could also be used to investigate the effects of varying oxygen availability on immune cell function against solid tumors.

In this work, a novel microfluidic TNBC immunoassay was developed to evaluate CAR-T cell-mediated cytotoxicity and target specificity against the EGF receptor using tumor cells co-cultured with stromal cells as spheroids. This is the first report showing a miniaturized assay for the assessment of chemotherapy, biologics and cell therapy studies in a 3D solid tumor platform. CAR-T cells rapidly targeted, disaggregated and killed cancer cells without significant cytotoxicity to the 3D stromal mass. This effect could be neutralized by blocking EGFR recognition by CAR-T. The cytotoxic effects of EGFR targeting CAR-Ts was demonstrated to be most pronounced when spheroids were pre-treated with chemotherapy in comparison to control and monotherapy conditions, likely as a result of chemotherapy induced upregulation of EGFR and PD-L1 expression by tumor cells. The platform enabled quantification of readouts, including on-target specificity.

The developed 3D microfluidic immunoassay has the potential to be a powerful tool in the development of immunotherapeutics, as well as facilitating the investigation of 3D tumor-immune cell interactions. In future, the miniaturization yet medium-throughput capabilities of the platform could be used to investigate the efficacy and specificity of a wide range of adoptive cell transfer therapies in combination with numerous other anti-cancer agents to identify the optimum regiment for a specific patient.

## Acknowledgements

This work was funded by AMS Biotechnology Europe Ltd (industrial PhD studentship to K.P and M.Z) and internal funds by Strathclyde University to M.Z. and K.P.

## Author contributions

K.P. performed all cell assay experiments. K.P., S.P. and T.M. optimised microfluidic protocols. K.P., S.B.C. and M.Z designed the experiments. All authors contributed to the writing, reviewing and editing of the manuscript.

