## Supplementary Information for "Assessment of CAR-T mediated and targeted cytotoxicity in 3D microfluidic TBNC co-culture models for combination therapy"

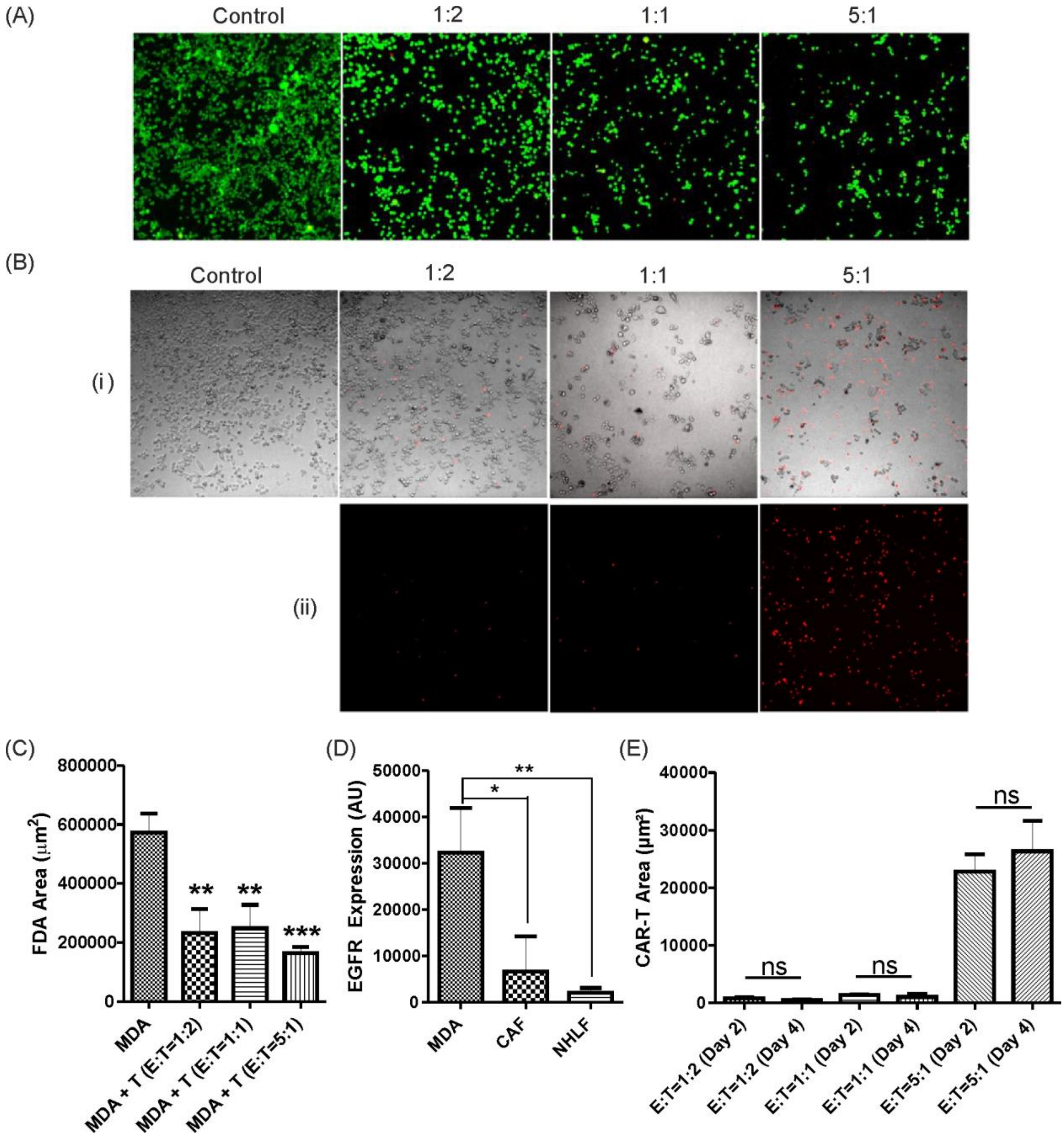


***Figure S1.*** *2D CAR-T assays. CAR-T cells were incubated with MDA-MB-468 cells for 72h at 5:1, 1:1 and 1:2 effector to target (E:T) ratios. (A) Representative viability staining with FDA (green) and PI (red) of MDA-MB-468 cells after 72h CAR-T incubation in a 96-well plate. (B) Representative (i) brightfield and (ii) fluorescence images of MDA-MB-468 (unlabelled) in a 96 well-plate, prior to removal of CAR-T cells (red). (C) Viability plot showing area of viable cancer cells after 72h CAR-T incubation (n=3). FDA and PI intensity values were normalized for all images. (D) Quantification of EGFR signal from MDA-MB-468, CAF and NHLF in 96-well plates after 3 days of culture and prior to CAR-T cell injection. (E) Plot showing area of labelled CAR-T cells after injection on day 2 of culture and after 48 hours of incubation with MDA-MB-468 in the different E:T conditions tested.*

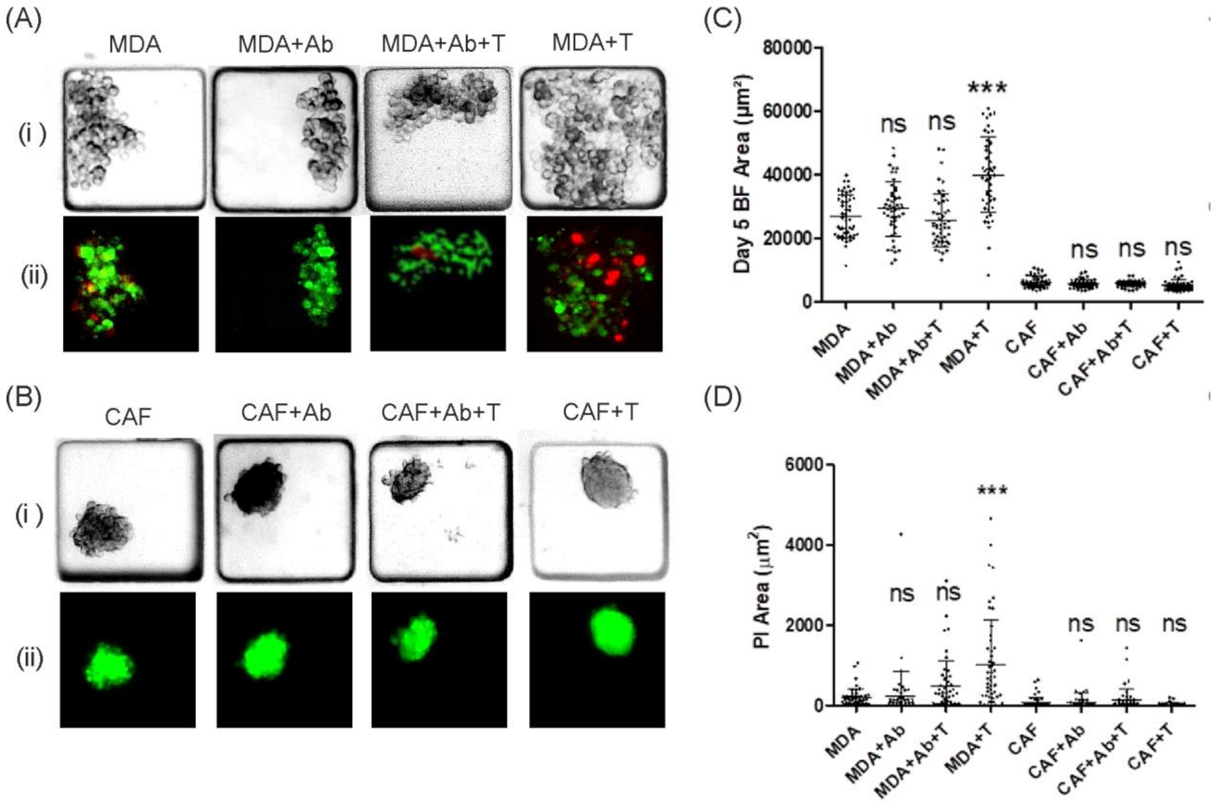


***Figure S2.*** *EGFR blocking assays in microfluidic spheroid cultures. (A) Representative (i) brightfield and (ii) fluorescent images of MDA-MDB-468 spheroid monocultures after 72h incubation with CAR-Ts, stained with FDA (green) and PI (red). (B) Representative (i) brightfield and (ii) fluorescent images of CAF (green) spheroid monocultures after 72h incubation with CAR-Ts, stained with PI (red). (C) Plot of spheroid area after CAR-T incubation (n=50). (D) Plot of PI signal area after CAR-T incubation (n=50).*

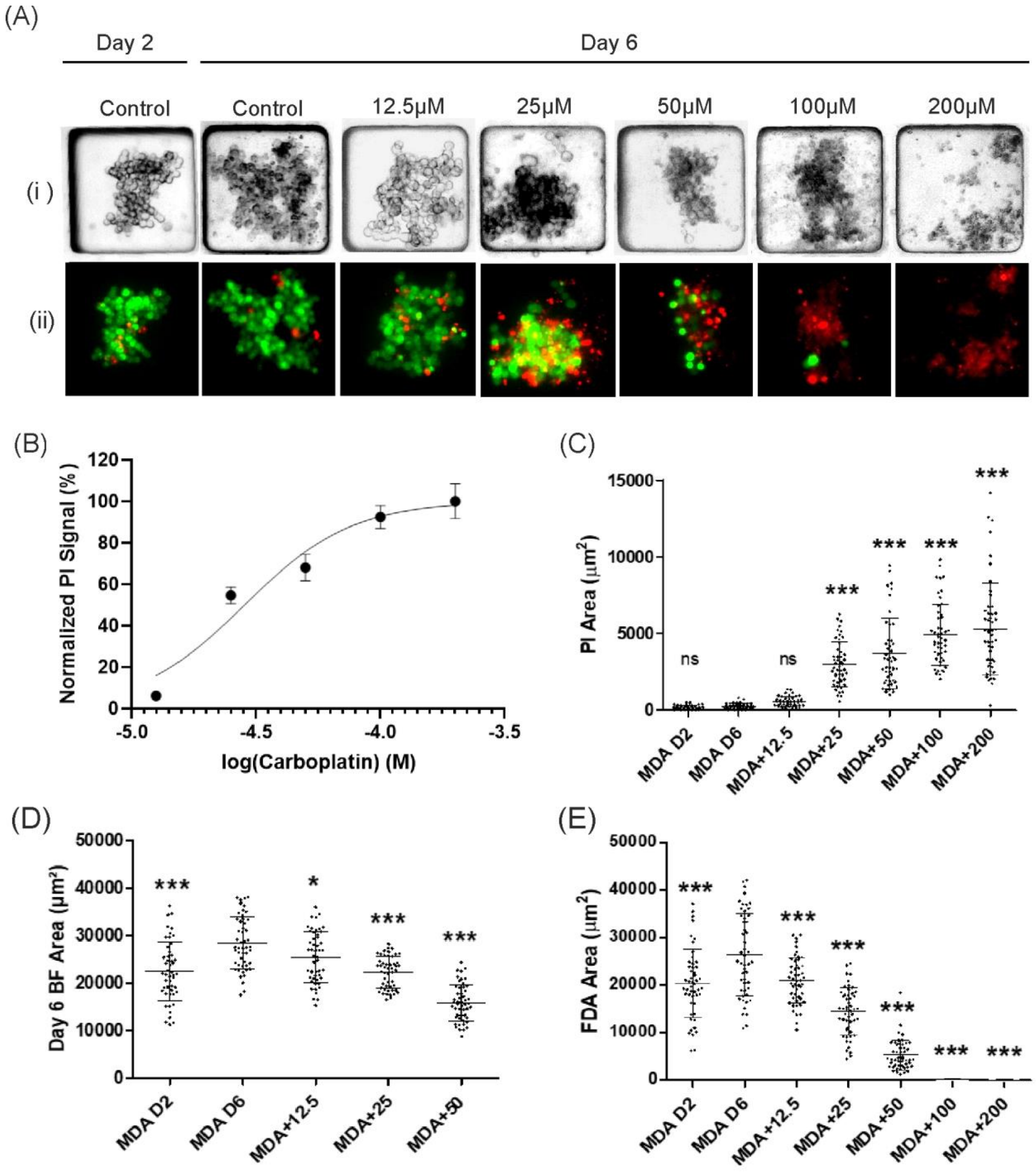


***Figure S3.*** *Effect of carboplatin treatment on tumour spheroid viability. (A) Representative (i) brightfield and (ii) fluorescent images of MDA-MDB-468 spheroid monocultures after formation (on day 2) and after exposure to carboplatin for 24h, staining with FDA (green) and PI (red) on day 6 of culture (endpoint). (B) Plot of normalized PI signal against log carboplatin concentration with error bars showing standard error of the mean (n=50). (C) Plot of PI signal area on day 6 (endpoint) of culture (n=50). (D) Plot of spheroid area from brightfield, on day 6 of culture (endpoint) (n=50). (E) Plot of viable spheroid area (from FDA signal) on day 6 (endpoint) of culture (n=50).*
